# Heterologous expression of the human cohesin complex in *Saccharomyces cerevisiae* results in a dominant-negative phenotype

**DOI:** 10.64898/2026.04.03.716359

**Authors:** Elizabeth Stephens, Akil Hamza, Maureen R. M. Driessen, Nigel J. O’Neil, Peter C. Stirling, Philip Hieter

## Abstract

The cohesin complex has conserved roles in sister chromatid cohesion, DNA replication, genome organization, and the DNA damage response. We heterologously expressed the human cohesin complex in yeast to probe the behaviour of human cohesin. Human cohesin was unable to complement loss of function mutations in yeast cohesin, either as single subunits or as complexes, including in the context of co-expressing up to 12 human cohesin-associated genes. Heterologous expression of human cohesin in yeast expressing wildtype yeast cohesin resulted in dominant cohesion dysregulation and DNA damage sensitivity phenotypes. We used co-immunoprecipitation to demonstrate that human SMC proteins interact with endogenous yeast cohesin rings creating dominant-negative hybrid complexes that disrupt endogenous cohesin biology.

## INTRODUCTION

The yeast genome is made up of approximately 6000 genes, of which more than 2000 have human orthologs (Kim et al., 2020). Over 700 human genes have been found to functionally complement the deletion of the corresponding yeast ortholog, many of which have implications for human disease (Kachroo et al., 2022). It is possible to use these “humanized” yeast models to study the genes and pathways associated with diseases, such as cancer, in an experimentally tractable model system. Humanized yeast provides a simple, high-throughput system to interrogate protein function, parse genotype-phenotype associations, characterize disease variants, and test small molecule inhibitors of human proteins (Hamza et al., 2015, 2020; Kachroo et al., 2022). Multigene complementation may be necessary to complement protein complexes. Although more complex than single gene complementation, it is possible to functionally replace entire yeast pathways and multi-protein complexes with their human counterparts (Boonekamp et al., 2022; Garge et al., 2020; Truong & Boeke, 2017; Agmon et al., 2020).

The cohesin complex is well conserved across species. Cohesin is a multifunctional complex made up of core subunits named SMC1A, SMC3, RAD21 and STAG1/2 in human and Smc1, Smc3, Mcd1, and Irr1 in *Saccharomyces cerevisiae* (**FIGURE 1A**). The cohesin complex has roles in sister chromatid cohesion, DNA damage repair, chromosome segregation and replication (Nasmyth & Haering, 2009). Cohesin also plays key roles in human genome organization and the maintenance of topologically associated domains (TADs) (Gassler et al., 2017; Merkenschlager & Nora, 2016; Rao et al., 2017; Schwarzer et al., 2017; Wutz et al., 2017). Cohesin mutations are common in a variety of cancers and are responsible for a group of disorders called cohesinopathies (Barbero, 2013; Di Nardo et al., 2022; Scott et al., 2025). Human cohesin is comprised of essential genes, making it difficult to study *in vivo*. A humanized yeast model could provide a platform to study cohesin mutations to better understand human cohesin biology and its role in disease.

**Figure 1:**
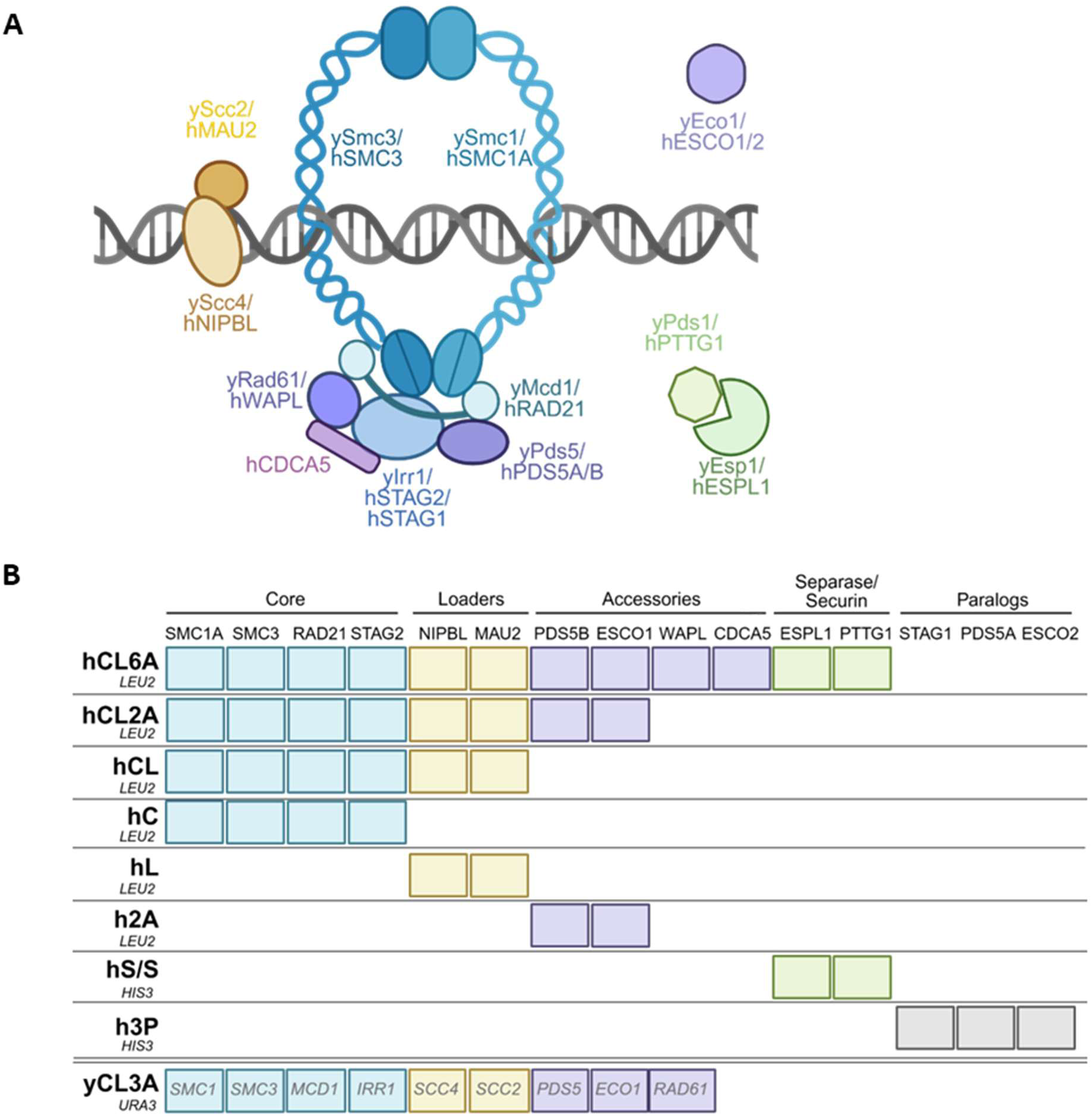
The cohesin complex and its partners. (A) Schematic of cohesin and its interacting partners highlighting the 4 core subunits, 2 loaders, and accessory proteins. (Created in BioRender. Fournier, L. (2026) https://BioRender.com/pc66ncw) (B) Schematic highlighting the various human or yeast cohesin gene combinations cloned on yeast plasmids or constructed by DNA synthesis (to create neochromosomes). *URA3*-marked yCL3A contains yeast cohesin genes flanked by endogenous promoters and heterologous terminators. The remaining plasmids (hCL6A, hCL2A, hCL, hC, hL, h2A, hS/S, and h3P) contain human cohesin genes flanked by the endogenous yeast promoter and terminator of the corresponding yeast ortholog (except for *CDCA5* which has no known yeast homolog, and was designed with the yeast promoter of *RPL13A* and terminator of *RPL22B*).

Here we report our attempts to replace the yeast cohesin complex with human cohesin using a systematic complementation strategy involving single gene and multi-gene replacement combinations. Heterologous expression of human cohesin was unable to complement the loss of yeast cohesin. Instead, we observed a dominant-negative phenotype consistent with dysfunctional cohesin. We found that the dominant-negative phenotype was the result of cross-species human-yeast cohesin complexes that associate with yeast chromatin and impact cohesin-associated phenotypes.

## RESULTS

### Human cohesin does not complement yeast cohesin, as single subunits or as a complex

To assess whether human cohesin could complement loss of yeast cohesin, we used temperature sensitive (Ts) mutants of the cohesin core components (*smc1-259*, *irr1-1*, *mcd1-73*), and the cohesin regulatory genes (*eco1-1, and pds1-128*). The human ortholog cDNAs (*SMC1A, STAG2, RAD21, ESCO1, PTTG1*) corresponding to each of the yeast cohesin genes were cloned into a yeast expression vector under the control of the GPD constitutive promoter. The transformed Ts strains were serially spotted and incubated at 25°C, 30°C and 37°C to assess complementation at the restrictive temperature (**SUPPLEMENTAL FIGURE 1A**). In all cases, the human cohesin orthologs did not rescue viability of the corresponding yeast Ts strains at the restrictive temperature. Notably, GPD promoter-driven *SMC1A* could not be transformed into yeast. To test whether constitutive expression of human SMC1A was deleterious in yeast, *SMC1A* was cloned into a galactose-inducible high-expression plasmid and transformed into wildtype and *smc1-259* strains in glucose media (**SUPPLEMENTAL FIGURE 1B**). Galactose inducible expression of *SMC1A* severely affected the growth of the wildtype strain and was synthetic lethal in the *smc1-259* background at permissive temperatures. This suggested that overexpression of human *SMC1A* in yeast resulted in a growth defect.

Human gene complementation studies in yeast have shown that some genes only complement when expressed together with their human protein interaction partners (Boonekamp et al., 2022; Garge et al., 2020; Truong & Boeke, 2017). To test whether co-expression of multiple human cohesin subunits or accessory factors could complement the Ts mutations in individual yeast cohesin subunits, we created neochromosomes (synthetic chromosomes) containing combinations of yeast codon optimized human cohesin and cohesin accessory genes under the control of the endogenous yeast promoters and terminators of the corresponding yeast orthologs (**FIGURE 1B**). The human cohesin neochromosomes included hC (4 core subunits), hCL (4 core + 2 loader subunits), hCL2A (4 core + 2 loader + 2 accessory subunits) and hCL6A (4 core + 2 loader + 6 accessory subunits). The difference between hCL2A and hCL6A is the addition of 4 human genes: one human gene that has nonessential yeast ortholog (*WAPL*), one human gene that has no known homolog in yeast (*CDCA5*), and two human genes that form a separate two-subunit complex composed of separase (*ESPL1*) and its inhibitor securin (*PTTG1*). To allow flexibility in setting up complementation assays with multiple combinations of human genes, we designed additional neochromosomes including hL (2 loader subunits), h2A (2 accessory subunits), hS/S (2 accessory subunits: separase and securin), and h3P (3 human paralogs). Since three human cohesin genes have multiple paralogs, we included the least diverged homologs (*STAG2*, *PDS5B*, *ESCO1*) in neochromosomes hC, hCL, hCL2A and hCL6A, but designed the separate h3P to include *STAG1*, *PDS5A* and *ESCO2*.

We first tested whether expression of the four human core (hC) subunits could complement Ts mutations in single yeast cohesin complex members (*smc1-259*, *smc3-1*, *mcd1-73* or *irr1-1*) by shifting growth to a non-permissive temperature. Expression of the human core complex did not complement single yeast mutants at non-permissive temperatures and resulted in synthetic growth defects in the cohesin mutants *smc3-1* and *mcd1-73* at permissive temperatures (**FIGURE 2A**). We also tested whether expression of the human cohesin loaders and human cohesin accessory genes together with the human core could complement the individual yeast Ts strains. All cohesin neochromosome combinations failed to complement single yeast cohesin Ts mutants at restrictive temperatures (**SUPPLEMENTAL FIGURE 2**).

**Figure 2:**
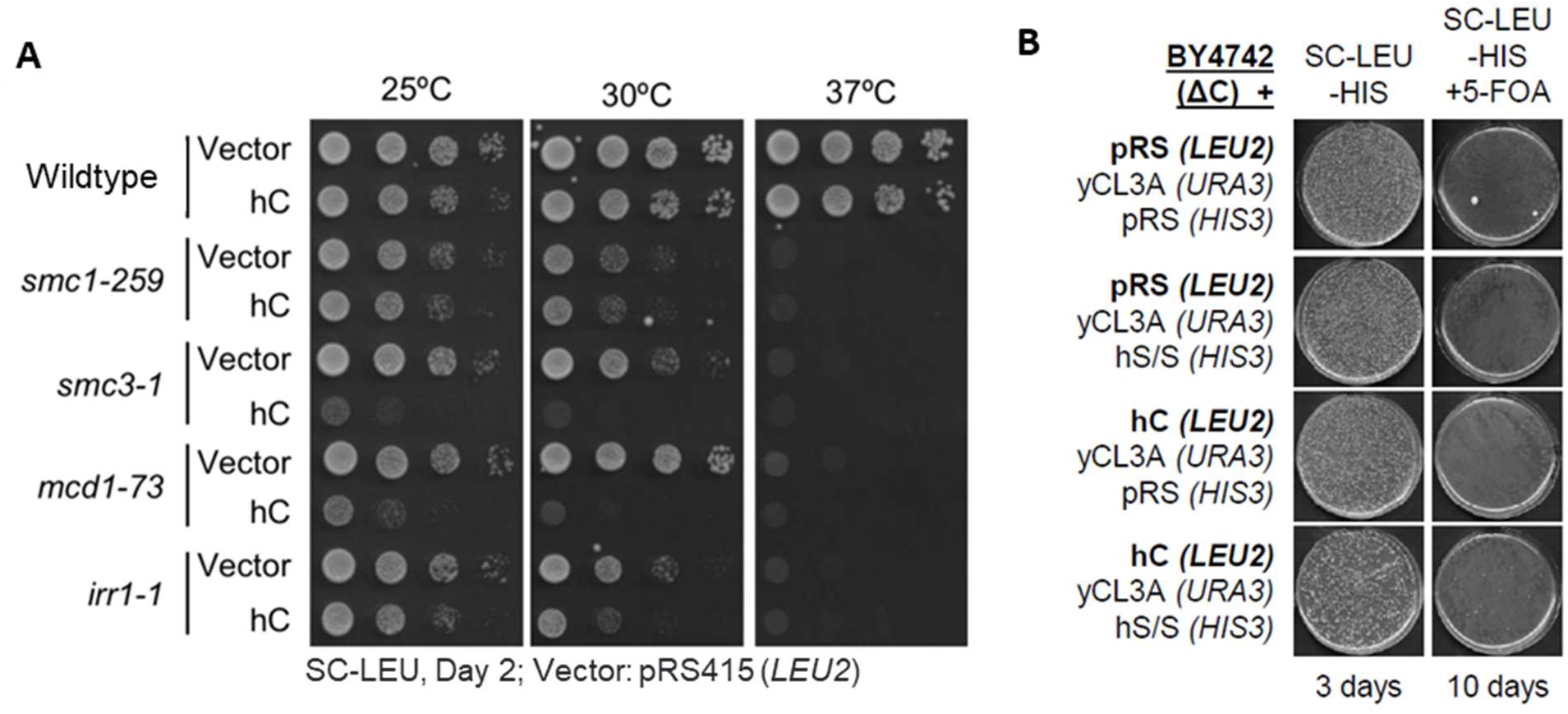
Human cohesin cannot functionally complement loss of yeast cohesin single subunits, or entire cohesin complex deletion mutants. (A) Heterologous expression of the four human cohesin core subunits (hC) does not complement loss of single yeast subunits at the restrictive temperature. (B) hC does not rescue viability of a ΔC yeast strain +/- hS/S. The yeast knockout strain (ΔC = *smc1Δ, smc3Δ, mcd1Δ, irr1Δ*) covered by the *URA3*-marked vector (yCL3A) was transformed with the indicated *LEU2*- and *HIS3*-marked vectors or recombinant constructs and maintained on −Ura −Leu −His media. Strains were plated on −Leu −His +5-FOA media to test complementation using 5-FOA plasmid shuffling. In the presented example, yeast strains were plated 1000-fold higher on 5-FOA plates compared to −Leu −His plates. Plates were incubated for 3-10 days at 30°C, with no complementation observed.

It is possible that residual yeast subunits could impair the complementation by human cohesin subunits. To test whether the entire yeast cohesin complex needed to be removed to allow for human complementation, a neochromosome marked by the counter-selectable *URA3* marker was generated to contain nine yeast cohesin genes orthologous to those on the human neochromosomes (yCL3A: 4 core + 2 loader + 3 accessory subunits) (**FIGURE 1B**). The yeast cohesin genes on the neochromosome were designed to be flanked by the yeast endogenous promoters and a set of heterologous terminators. The heterologous terminators were introduced to allow specific knockout of the endogenous genomic ORFs by using CRISPR to target the endogenous terminators in the genome. We confirmed that the yeast neochromosome yCL3A can complement cohesin Ts alleles at restrictive temperatures (**SUPPLEMENTAL FIGURE 3A**). We constructed a series of strains with multiple genomic deletions of yeast cohesin genes in the presence of yCL3A (ΔC, ΔCL, ΔCL2A, ΔCL3A). The yeast neochromosome complemented the simultaneous deletion of all nine genome cohesin genes from the genome (**SUPPLEMENTAL FIGURE 3B**). To test if human cohesin could complement simultaneous loss of yeast cohesin subunits, the human neochromosomes were transformed into cells containing a yeast neochromosome complementing the genomic deletions of cohesin and cohesin associated genes. Dual neochromosome containing cells were plated onto media containing 5-FOA, which counter-selects against the *URA3* marker on the neochromosome containing the yeast cohesin genes. This plasmid swap approach was used to test for complementation by hC, hCL, and hCL2A of the simultaneous deletions of the corresponding yeast genes. To eliminate the possibility that a determent to complementation is the inability of human core cohesin proteins to disengage from yeast DNA without a species-specific separase, we included hS/S in different combinations with the other human cohesin neochromosomes. When complementation was not observed, we introduced up to 15 human cohesin genes (hCL6A and h3P) into a yeast strain that had genomic deletions of 9 yeast cohesin genes. None of the multigene combinations complemented loss of the cohesin complex (**FIGURE 2B**, **SUPPLEMENTAL FIGURE 3B**) (Summarized in **SUPPLEMENTAL TABLE 1**).

### Expression of human SMC1A or SMC3 causes a dominant-negative phenotype

Although human cohesin did not complement loss of yeast cohesin, we observed that expression of human cohesin caused a growth defect in wildtype yeast cells. Expression of human cohesin (hC) also caused growth defects in *smc3-1* and *mcd1-73* cells at permissive temperatures (**FIGURE 2A**). To determine if the phenotype was exacerbated by the addition of human cohesin regulatory subunits, which may have an impact on the ability of human core subunits to load on yeast chromatin, we co-expressed the human core cohesin genes with human cohesin loader and accessory genes (hCL, hCL2A, hCL6A). Co-expression of the human cohesin core and cohesin loaders enhanced the growth defects of human cohesin in wildtype cells (**FIGURE 3A**). In contrast, expression of human cohesin loader genes (hL) or accessory genes (h2A) alone did not result in a growth defect. The fact that the human core subunits cause a fitness defect that is exacerbated by addition of the human cohesin loaders, suggests that the phenotype may result from human cohesin subunits loading onto yeast DNA resulting in a dominant-negative effect as described by Herskowitz (1987). To test if human cohesin was loading onto chromatin, we performed chromatin fractionation analysis and found that the 4 human core subunits are stably expressed and are found in both the cytoplasmic and chromatin fractions (**FIGURE 3B**). We next tested whether the dominant-negative phenotype could be suppressed by co-expressing human separase and securin (hS/S) together with the human core cohesin. Expression of human separase and securin did not suppress the dominant-negative effect of heterologous expression of human cohesin core subunits in yeast (**SUPPLEMENTAL FIGURE 4B**).

**Figure 3:**
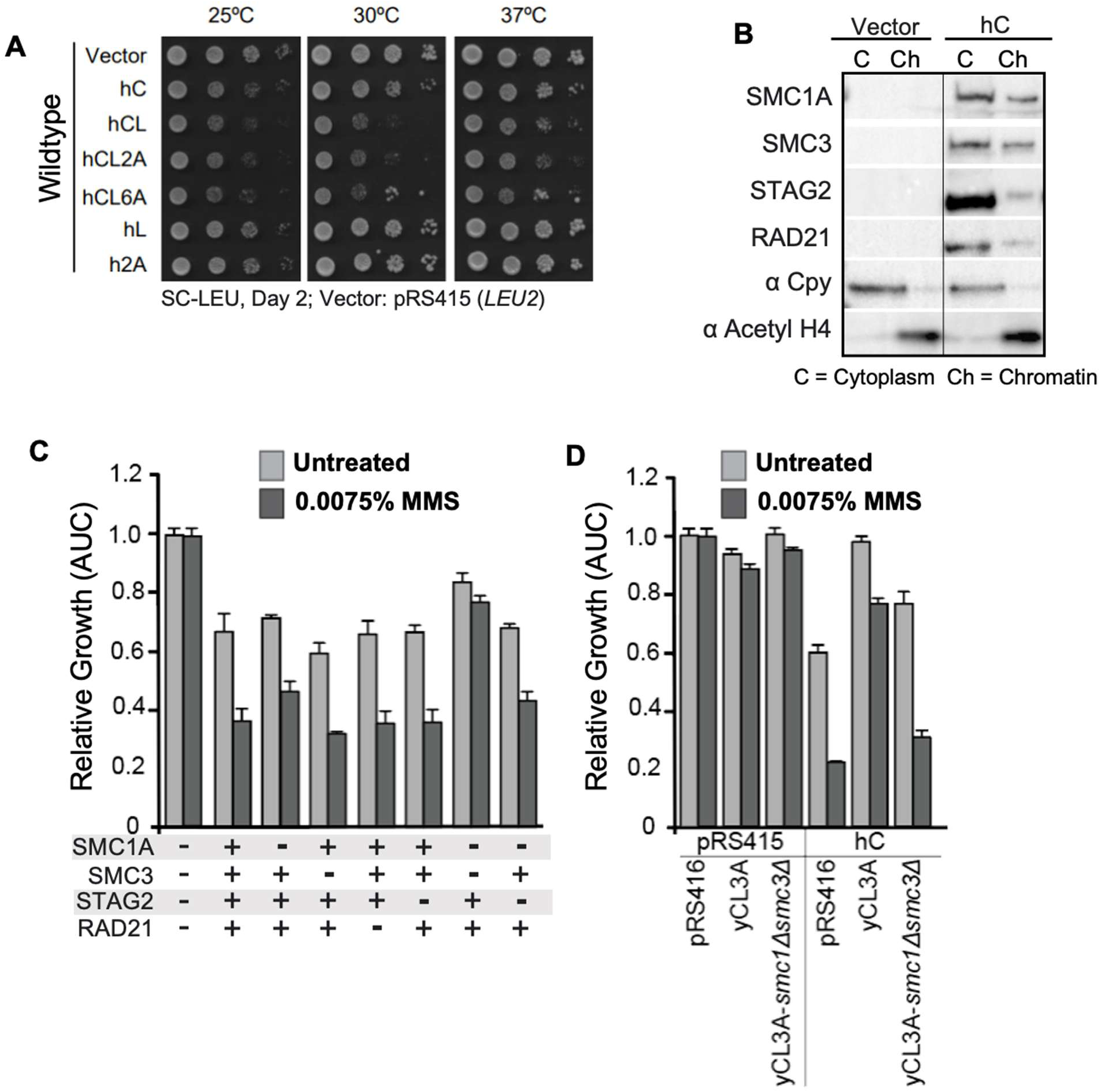
The SMC1A and SMC3 subunits cause a dominant-negative phenotype in cells containing endogenous yeast cohesin, and this effect is suppressed with extra copies of Smc1 and Smc3. (A) Growth assay showing that expression of human cohesin core proteins from hC, hCL, hCL2A, and hCL6A cause a dominant phenotype in wildtype yeast, whereas hL and h2A alone have no effect on growth. The dominant negative growth phenotype is stronger when the human core and loader complex are expressed at the same time (hCL, hCL2A, hCL6A). (B) Chromatin fractionation showing that human cohesin core proteins expressed from hC are found in the chromatin fraction, suggesting interaction of the human subunits with yeast chromatin (C= cytoplasmic faction, Ch= Chromatin fraction). αCpy and αAcH4 are controls indicating successful fractionation. (C) Growth curve analysis of systematic deletions of proteins from the hC showing that presence of the SMC proteins cause a dominant sensitivity to MMS (0.0075%) and this can only be rescued by the deletion of both *SMC1A* and *SMC3* simultaneously. (D) Growth curve analysis showing that rescue of dominant-negative growth phenotypes of yeast containing hC by yCL3A is dependent on yeast Smc1 and Smc3. For C-D liquid growth assays, each strain was tested in three replicates per condition and area under the curve (AUC) value was calculated for each replicate. Strain fitness was defined as the AUC of each yeast strain relative to the AUC of the wildtype strain (BY4742) containing the vector control and grown in the same media condition (mean +/- SD).

If human cohesin is associating with yeast chromatin, it may disrupt the function of yeast cohesin causing other phenotypes associated with cohesion dysfunction. At permissive temperatures, cohesin Ts mutants are sensitive to the DNA-damaging agent methyl methanesulfonate (MMS) (Svensson et al., 2011). We therefore measured the effects of expressing human cohesin genes in wildtype cells in response to low doses of MMS. Quantitative liquid growth curves confirmed that expression of the human core cohesin genes resulted in a growth defect and that the growth defect was exacerbated by the addition of a low concentration of MMS (**FIGURE 3C**).

To better understand how heterologously expressed human cohesin was causing a dominant-negative effect, we deleted each of the four core genes from the hC neochromosome, individually and in combination. We measured the effect on sensitivity to MMS. Single deletions of any of the four core genes had no impact on MMS sensitivity, however simultaneous deletion of both *SMC1A* and *SMC3* ameliorated the sensitivity to MMS. This suggests that expression of either human SMC cohesin subunit can cause a dominant-negative phenotype (**FIGURE 3C**).

Dominant-negative effects can be the result of incorporation of mutant dysfunctional proteoforms in complexes, which can poison the activity of the complex (Herskowitz, 1987). To determine if the dominant-negative effect of heterologous expression of human cohesin could be rescued by overexpressing yeast cohesin subunits, yeast genes (*SMC1*, *SMC3*, *IRR1*, *SCC4*, *PDS5*, *ECO1* or *RAD61*) were each singly overexpressed from a galactose-inducible promoter. Growth was measured in the presence and absence of a low concentration of MMS (0.0075%). When co-expressed with the hC neochromosome, overexpression of single yeast cohesin subunits did not suppress the hC dominant-negative phenotype (**SUPPLEMENTAL FIGURE 4A**). To assess if overexpression of multiple yeast genes can suppress of the dominant-negative phenotype, we introduced the endogenously-regulated yCL3A into wildtype yeast to supplement the endogenous genomic cohesin genes with a second copy. This increased gene dosage suppressed the dominant-negative phenotype of hC, both in the presence and absence of MMS (**SUPPLEMENTAL FIGURE 4B**). To determine which yeast subunits were responsible for rescuing the dominant-negative effect caused by human *SMC1A* and *SMC3*, we deleted yeast *SMC1* and *SMC3* from the yCL3A neochromosome and showed that the suppression of the dominant-negative phenotype of human cohesin expression was abolished (**FIGURE 3D**). This suggests that both Smc1 and Smc3 are needed to outcompete SMC1A and SMC3 to mitigate the dominant-negative phenotype effect of heterologous expression of human SMC genes.

### Dominant-negative phenotype of heterologously expressed human SMC proteins may be due to interaction with yeast subunits

Given that the dominant-negative phenotype did not require the entire human cohesin core, we hypothesized that human cohesin subunits may be forming hybrid cohesin complexes that contain both human and yeast subunits and that these hybrid cohesin complexes were dysfunctional. To test whether human cohesin loading was dependent on SMC1A and SMC3 proteins, we performed chromatin fractionation with systematic deletions of the cohesin subunit genes from the hC neochromosome (hC, hC-*smc1aΔ*, hC-*smc3Δ*, hC-*smc1aΔ smc3Δ*, hC-*stag2Δ* and hC-*rad21Δ*). When *SMC1A* or *SMC3* are deleted individually, the other human cohesin subunits are still associated with the yeast chromatin fraction (**FIGURE 4A**). When *RAD21* is deleted, SMC1A and SMC3 are still associated with yeast chromatin but STAG2 is greatly reduced, consistent with STAG2 directly associating with RAD21 (Deardorff et al., 2012; Nasmyth & Haering, 2009). Conversely, deletion of *STAG2* did not affect RAD21 association with yeast chromatin. When both human SMC subunits were deleted, none of the human cohesin subunits were present on yeast chromatin demonstrating that chromatin loading of RAD21 and STAG2 subunits requires at least one human SMC protein to be present. This suggests that human cohesin subunits are loading onto yeast DNA in the absence of a complete human core and that this association requires at least one human SMC cohesin subunit.

**Figure 4:**
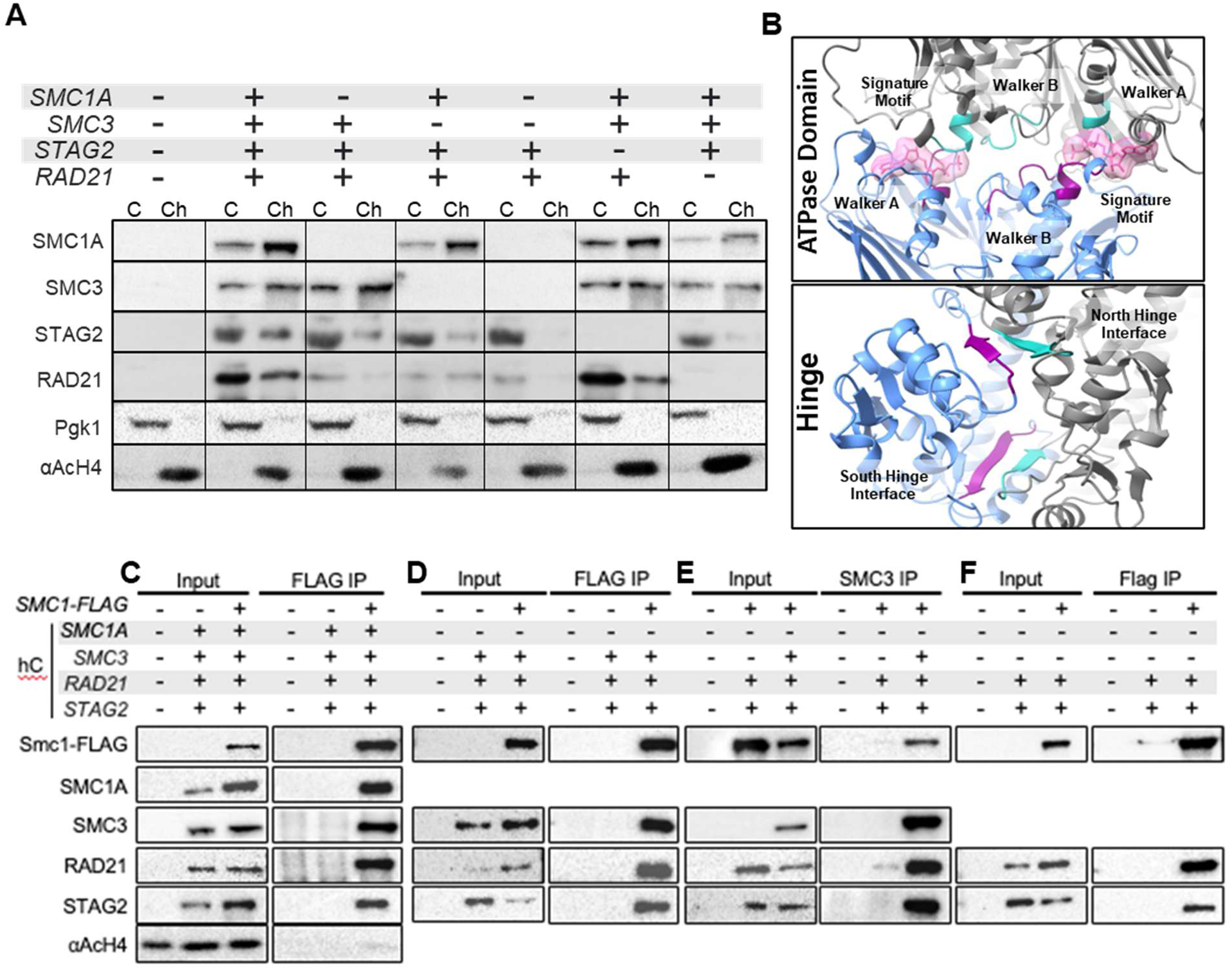
Yeast and human cohesin subunits form hybrid complexes. (A) Chromatin fractionation showing that the human core subunits are found in the chromatin fraction. Systematic deletion of hC subunits show that the core subunits are associating with yeast chromatin even when a complete human core complex cannot be formed, suggesting yeast-human protein interactions. Pgk1 and αAcH4 are controls indicating successful fractionation (C= cytoplasmic faction, Ch= Chromatin fraction). (B) AlphaFold3 predictions showing that yeast Smc3 (grey) and human SMC1A (blue) are predicted to dimerize and form an ATPase domain that can bind ATP (pink) and interact via their hinge domains. (C) Co-IP analysis shows pulldown of SMC1A and SMC3 when Smc1-FLAG is immunoprecipitated. (D) In the absence of SMC1A (hC-1Δ) SMC3 co-immunoprecipitated with Smc1-FLAG, confirming human-yeast hybrids. (E) Immunoprecipitation of SMC3 shows pulldown of Smc1-FLAG, confirming this interaction. (F) When Smc1-FLAG is immunoprecipitated in the presence of hC-*smc1aΔ smc3Δ* (hC-1/3Δ), both RAD21 and STAG2 co-immunoprecipitated.

To determine if the interaction interfaces between human and yeast cohesin subunits were potentially compatible, we used Alphafold3 to model interactions between yeast and human cohesin subunits. Human SMC1A and yeast Smc3 are predicted to be compatible with each other at both their hinge domains and ATP binding heads (**FIGURE 4B**). To test if yeast Smc1 was interacting with human SMC proteins, we expressed human core cohesin in a strain ectopically expressing Smc1 tagged with FLAG to facilitate immunoprecipitation assays. Both human SMC3 and SMC1A co-immunoprecipitated with Smc1-FLAG (**FIGURE 4C**). The co-immunoprecipitation of human SMC1A with yeast Smc1-FLAG was not surprising as cohesin SMCs can co-immunoprecipitate with themselves (Zhang et al., 2008). It is possible that the complete human cohesin complexes were co-immunoprecipitating with Smc1-FLAG. To test whether hybrid yeast human cohesin complexes were forming, we performed a co-IP using Smc1-FLAG in a strain carrying a human neochromosome lacking *SMC1A* (hC-*smc1aΔ*) to prevent the potential formation of complete human cohesin rings on yeast chromatin. SMC3, RAD21 and STAG2 co-immunoprecipitated with yeast Smc1-FLAG (**FIGURE 4D**). The reciprocal experiment was performed by immunoprecipitating SMC3. Western blot analyses showed that yeast Smc1-FLAG co-immunoprecipitated with human SMC3 (**FIGURE 4E**). Because RAD21 and STAG2 are not effectively loaded onto yeast DNA in the absence of both SMC1A and SMC3 subunits (**FIGURE 4A**), we wanted to evaluate if yeast Smc1 can interact with these human subunits. In a strain carrying hC-*smc1aΔsmc3Δ* and expressing Smc1-FLAG, co-IP shows that Smc1-FLAG is interacting with RAD21 and STAG2 (**FIGURE 4E**). This was surprising as expression of human RAD21 and STAG2 did not cause a dominant-negative phenotype (**FIGURE 3**) and they do not associate with yeast DNA in absence of human SMCs (**FIGURE 4A**). It is possible that these hybrid complexes may form but not load onto DNA, unlike hybrid complexes containing human SMCs.

No commercial antibodies were available to probe the other yeast cohesin subunits and it was not possible to tag all the cohesin subunits without affecting viability, so we used single-molecule protein sequencing (SMPS) with a Quantum-Si Platinum^®^ benchtop sequencer to confirm the presence of yeast cohesin subunits in the co-immunoprecipitation (Reed et al., 2022). To test the ability of the SMPS to identify cohesin subunits from co-immunoprecipitation, we first prepared peptide libraries from anti-FLAG pulldowns of yeast lysates containing the Smc1-FLAG but not expressing human cohesin subunits. SMPS identified 3,495 aligned reads that mapped to the yeast cohesin proteins (**TABLE 1; SUPPLEMENTAL FIGURE 5**). As expected, the most abundant protein was Smc1-FLAG with 2,700 aligned reads mapping to 28 predicted peptides (10 with a false discovery rate (FDR) < 0.1). The next most abundant was Smc3 with 489 aligned reads mapping to 28 predicted peptides (6 with an FDR <0.1). This demonstrated that this approach could identify yeast cohesin proteins from a co- immunoprecipitation experiment. We next co-immunoprecipitated human SMC3 with an anti-SMC3 antibody from cells expressing *SMC3*, *RAD21* and *STAG2* (but not *SMC1A*) and performed SMPS. This identified 215 reads aligning to 17 peptides from yeast Smc1 (3 peptides, FDR <0.1) and 222 reads aligning to 22 peptides from yeast Smc3 (4 FDR <0.1). Together, these data suggest that human SMC1A and SMC3 can interact with yeast Smc1 and Smc3 to form hybrid cohesin complexes. These hybrid complexes could be dysfunctional causing a dominant-negative phenotype.

**TABLE 1:**
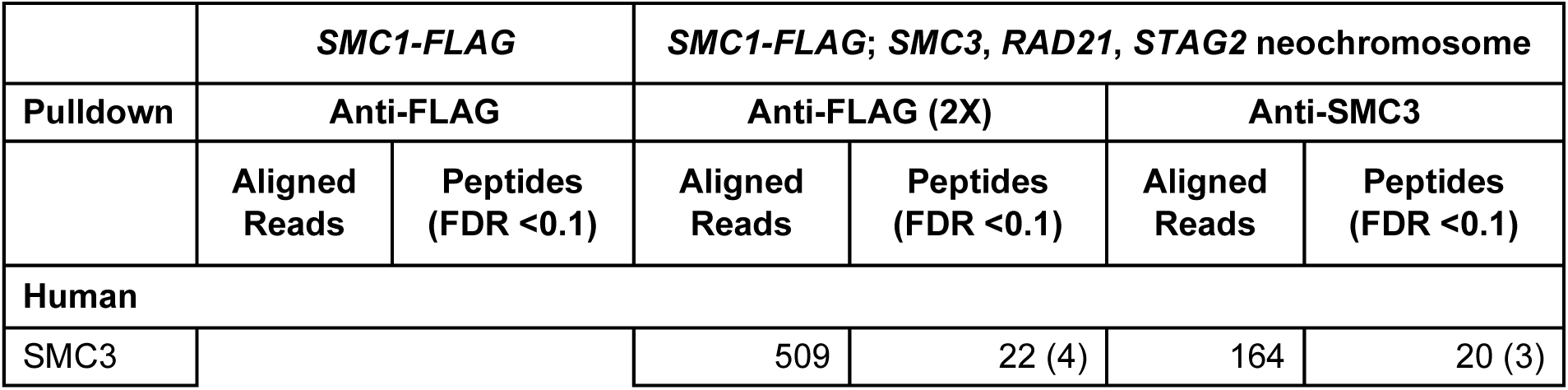
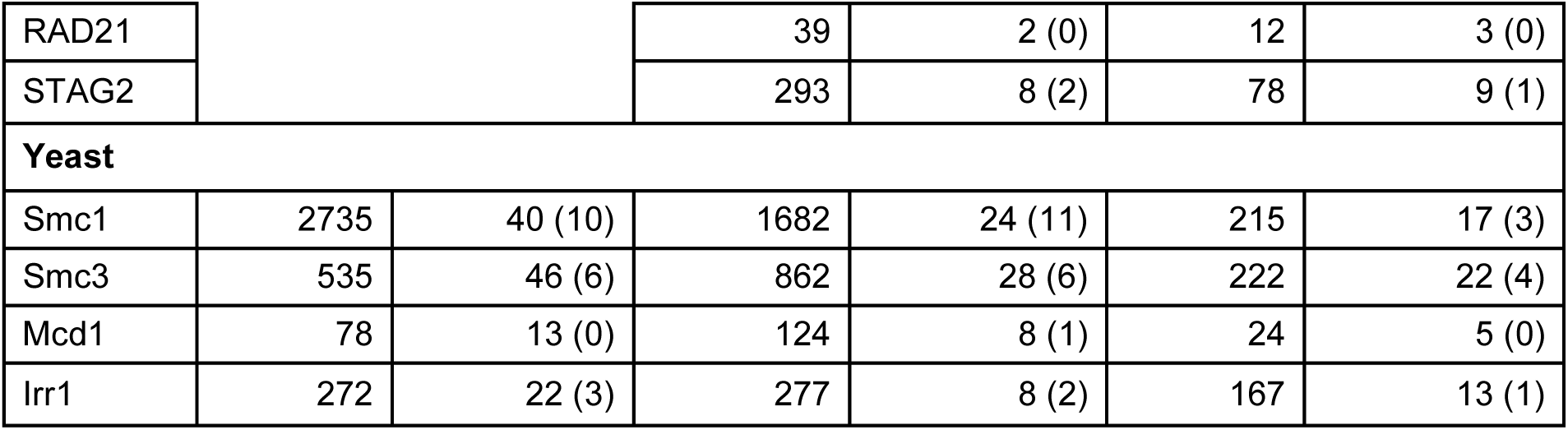
Peptides identified by SMPS from co-immunoprecipitation experiments.

|  | *SMC1-FLAG* |  | *SMC1-FLAG; SMC3, RAD21, STAG2* neochromosome |  |  |  |
| --- | --- | --- | --- | --- | --- | --- |
| Pulldown | Anti-FLAG |  | Anti-FLAG (2X) |  | Anti-SMC3 |  |
|  | Aligned Reads | Peptides (FDR <0.1) | Aligned Reads | Peptides (FDR <0.1) | Aligned Reads | Peptides (FDR <0.1) |
| **Human** |  |  |  |  |  |  |
| SMC3 |  |  | 509 | 22 (4) | 164 | 20 (3) |
| RAD21 |  |  | 39 | 2 (0) | 12 | 3 (0) |
| STAG2 |  |  | 293 | 8 (2) | 78 | 9 (1) |
| **Yeast** |  |  |  |  |  |  |
| Smc1 | 2735 | 40 (10) | 1682 | 24 (11) | 215 | 17 (3) |
| Smc3 | 535 | 46 (6) | 862 | 28 (6) | 222 | 22 (4) |
| Mcd1 | 78 | 13 (0) | 124 | 8 (1) | 24 | 5 (0) |
| Irr1 | 272 | 22 (3) | 277 | 8 (2) | 167 | 13 (1) |

### Expression of human cohesin leads to DNA repair centres and mitotic defects

To test which cellular processes in yeast are affected by the presence of human cohesin, we expressed human cohesin core genes in yeast lacking *RAD52*, which is needed for homologous recombination repair, or lacking *RAD61*, which promotes the removal of cohesin from chromatin, or lacking *CTF8*, which promotes replication fork restart and replication-coupled cohesion establishment. Heterologous expression of human cohesin was synthetic lethal in *rad61Δ* and *ctf8Δ* demonstrating that the presence of human cohesin core proteins exacerbated cohesin defects. Expression of human cohesin in *rad52Δ* resulted in slow growth, which suggests that human cohesin was causing DNA damage or replication stress that required homologous recombination for resolution. The synthetic lethality and growth defects were not observed when both *SMC1A* and *SMC3* genes were deleted from the hC neochromosome (**FIGURE 5A**).

**Figure 5:**
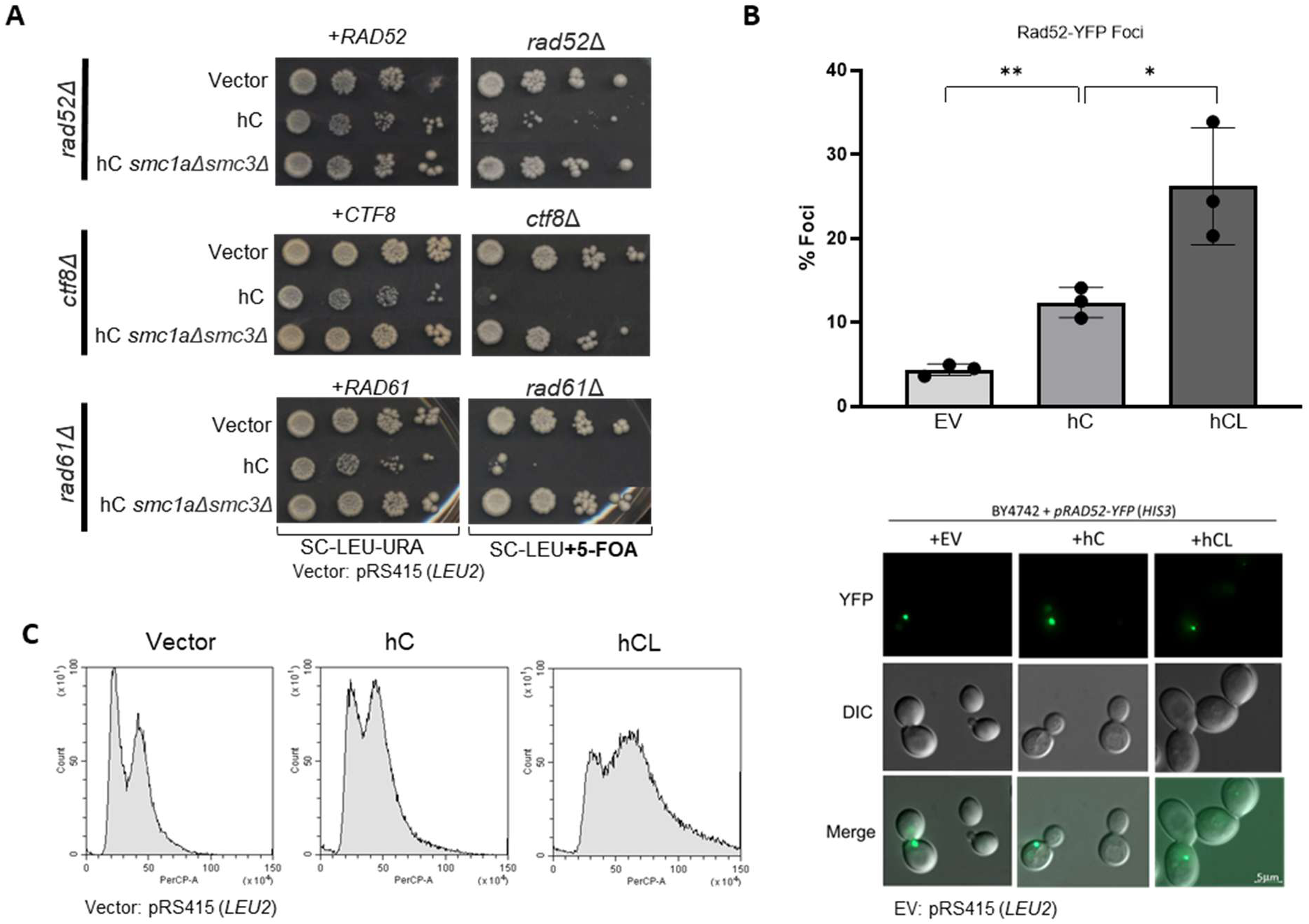
Human cohesin causes dominant DNA damage and mitotic defects. (A) Serial dilutions using a 5-FOA plasmid shuffle system shows synthetic lethal interactions between hC and *ctf8Δ* or *rad61Δ,* and synthetic slow growth with a *rad52Δ* background. Left panels show strains carrying the *LEU2*-marked hC and a *URA3*-marked plasmid containing a copy of the gene deleted genomically. In the right panels, cells are plated on media containing 5-FOA, uncovering the genomic deletion and revealing synthetic lethal interactions that are dependent on SMC1A and SMC3. (B) Rad52-YFP foci, assessed in a wildtype background, shows a dominant significant increase when hC proteins (SMC1A, SMC3, RAD21 and STAG2) are expressed (EV 5% vs hC 12%) and a further significant increase when the loaders (NIPBL and MAU2) are added (hC 12% vs hCL 24%). Error bars represent ± SD, n=3, >100 cells each. *, p<0.05, **, p<0.01, unpaired Student’s *t-*test. Scale bar: 5µm. (C) Cell cycle analysis profiles show that hC causes a moderate accumulation of cells in G2 vs empty vector, and hCL causes a shift to G2 and super G2 cells.

The genetic interaction with *rad52Δ* led us to evaluate whether expression of human cohesin was resulting in an increase in repair intermediates by assessing Rad52-YFP foci. Heterologous expression of human cohesin caused a significant increase in Rad52-YFP foci that was exacerbated when the human cohesin loaders were co-expressed with human cohesin (**FIGURE 5B**). While imaging Rad52-YFP foci, we noted that expression of human cohesin had effects on cellular morphology that was indicative of cells stalled in late S or G2 of the cell cycle. Flow cytometry was used to quantify the large-budded cells based on DNA content. Heterologous expression of human cohesin resulted in an increase in large-budded cells that was further exacerbated by co-expression of the human cohesin loaders, suggesting that expression and loading of human cohesin onto yeast chromatin was causing cells to accumulate in the late S or G2 phases (**FIGURE 5C**).

### Expression of mutant human SMC proteoforms does not mitigate or enhance the dominant-negative phenotype

There are SMC cohesin gene mutations that have specific functional effects. We hypothesized that the human-yeast hybrid complexes may not bind or hydrolyse ATP, which is necessary for cohesin function. We tested a series of SMC mutations: *smc1a-L1129V*, which enables cohesion without ATP hydrolysis or acetylation by ESCO1 (EcoI), *smc1a-K39I* and *smc3-K38I*, which cannot bind ATP, and *smc1a-E1158Q* and *smc3-E1144Q*, which can bind but not hydrolyze ATP (Arumugam et al., 2003; Elbatsh et al., 2016; Heidinger-Pauli et al., 2010; Marcos-Alcalde et al., 2017). Expression of these human SMC mutations did not enhance or suppress the chromatin association, phenotypic effect or synthetic lethality with *rad52Δ*, *rad61Δ* or *ctf8Δ* (**FIGURE 6A, B, C**). Since mutagenizing a single subunit may not show a measurable phenotype in growth and synthetic lethal interactions, we performed chromatin fractionations of the human cohesins expressing mutant SMC3 proteins (K38I or E1144Q). We would expect to see less protein in the chromatin fraction expressing the *smc3-K38I* mutation, as the lack of ATP binding blocks DNA loading (Arumugam et al., 2003). However, chromatin loading is not affected by expression of either the Smc3-K38I or the Smc3-E1144Q mutant protein. These data demonstrate that the dominant-negative phenotype from expression of human SMC is not dependent on ATP binding or hydrolysis.

**Figure 6:**
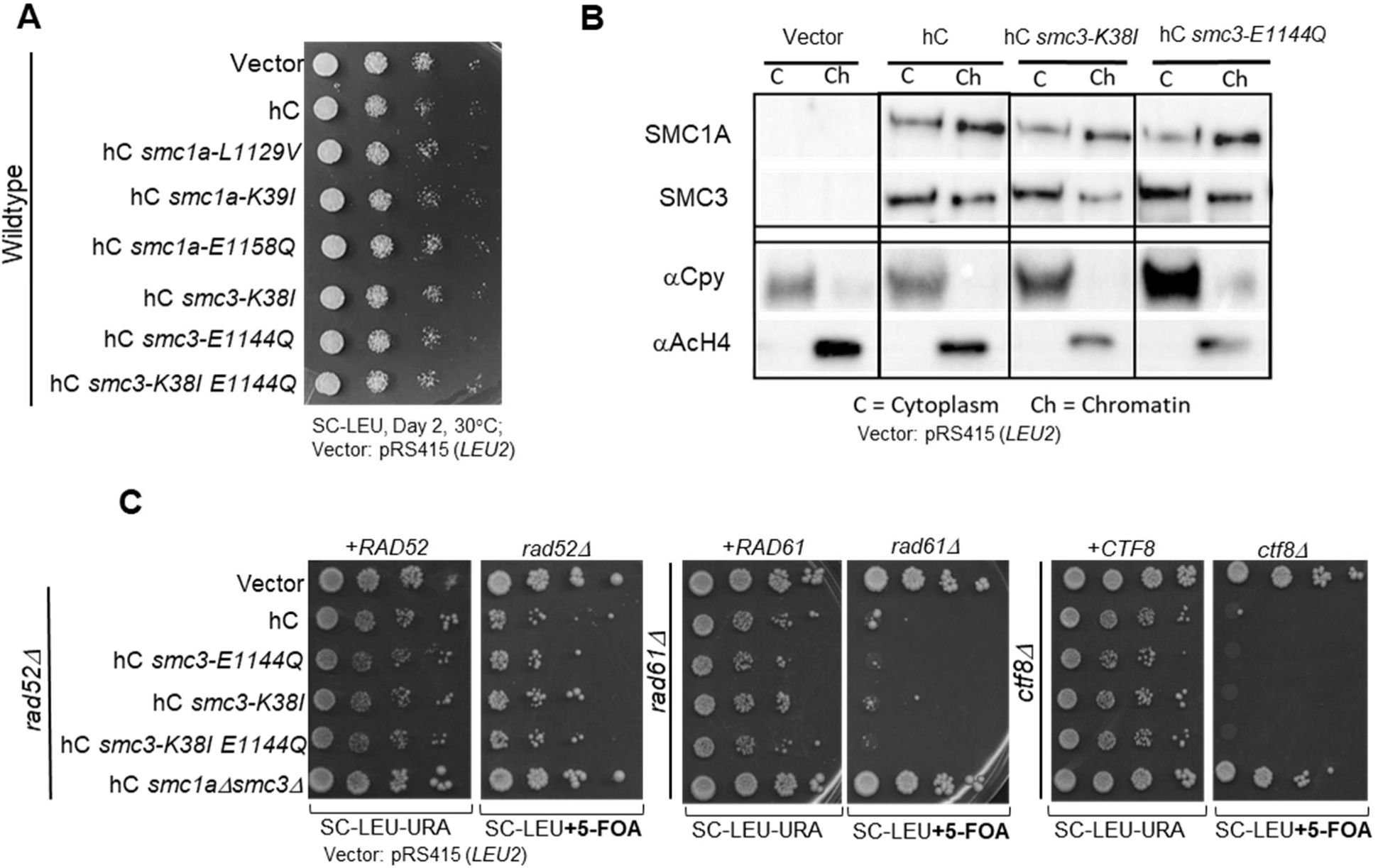
Cohesin-disrupting mutations do not ameliorate the dominant-negative phenotypes of hC. (A) Serial spot dilutions of wildtype yeast ectopically expressing hC, with cohesin-function altering *SMC1A* or *SMC3* mutations. Expression of neochromosomes containing mutated SMC proteins do not cause an altered growth phenotype. (B) Chromatin fractionation analysis assessing chromatin association of SMC3 harbouring *E1144Q* or *K38I mutations*. Expression of mutant SMC3 protein does not alter association of SMC3 with yeast chromatin. αCpy and αAcH4 are controls indicating successful fractionation (C= cytoplasmic faction, Ch= Chromatin fraction). (C) Synthetic lethal testing between hC-*smc3* mutants and *rad52Δ, rad61Δ* or *ctf8Δ* in plasmid shuffle strains. The mutant proteins do not alter the synthetic lethal phenotypes caused by hC.

## DISCUSSION

We attempted to functionally replace one to nine yeast cohesin complex subunits with human cohesin subunits singly or as an entire complex. Complementation studies have shown that some protein complexes and pathways may require co-expression of multiple genes simultaneously to achieve complementation (Davey et al., 2011; Gao et al., 2005; Paul et al., 2015). We constructed and tested neochromosomes containing increasingly complex combinations of two to fifteen genes encoding human cohesin subunits, loaders and accessory factors. However, even simultaneously replacing nine cohesin and cohesin-associated yeast genes with their human counterparts failed to achieve complementation. Several factors affect cross-species complementation including gene modularity and cellular role (Kachroo et al., 2015). Given the large number of genetic and physical interactions of cohesin and the diverse roles of cohesin, it was possible at the outset of this study that human cohesin components would be unable to functionally replace the corresponding yeast cohesin components. To date, few studies have reported successful multicomponent humanization, with the exceptions being the humanization of the nucleosome (Truong & Boeke, 2017), and the glycolysis pathway, which included the humanization of 25 genes (Boonekamp et al., 2022). In the case of the glycolysis pathway, it may have been successful because the pathway is a self-contained biochemical pathway dependent on substrates that are the same in both yeast and human cells. This is illustrated by the fact that the glycolysis pathway can be reconstituted entirely *in vitro* (Liu et al., 2017). Even though it could be reconstituted *in vitro*, the humanized pathway expressed in yeast did not support normal growth rates and this was explored using adaptive laboratory evolution. The authors were able to evolve a strain expressing the human glycolytic pathway that had much higher fitness demonstrating the need for the pathway to be adapted to the yeast cellular environment. Evolution was also needed for the humanized nucleosome to fully rescue loss of the yeast nucleosome, demonstrating the challenges of interspecies complementation of a multi-subunit complex (Truong & Boeke, 2017). Consistent with this added complexity, it is predicted that genes with many interacting partners also have a lower chance of complementation. In humans, the cohesin pathway has evolved to have an increasingly complex role with many different functions. Furthermore, human cohesin has multiple genes that have paralogs further complicating humanization efforts (Laurent et al., 2020). It may be possible to use adaptive laboratory evolution to generate an evolved human cohesin complex that does complement loss of yeast cohesin. We have shown that human cohesin interacts with its yeast counterparts. Given the complexity of evolving the entire human cohesin ring simultaneously, it might be easier to attempt to evolve a single human cohesin subunit so that it is functional in a cohesin ring with yeast subunits. In this way, each human cohesin subunit could be optimized to function in the yeast cellular environment.

Heterologous expression of human genes can be informative even if they do not complement loss-of-function mutations in their corresponding yeast orthologs. For example, heterologous expression of *PARP1* and *PARP2*, which lack a yeast homolog, was used to discover new small molecule inhibitors (Perkins et al., 2001). Similarly, the Alzheimer’s disease associated beta-amyloid peptide was expressed in yeast to find factors affecting disease biology and risk factors (Treusch et al., 2011). Although, expression of human cohesin did not complement loss of yeast cohesin, human cohesin appears to load onto yeast chromatin and result in dominant-negative phenotypes that are associated with the biological roles of cohesin; response to DNA damage, cell cycle progression, and mitosis. This suggests that ectopic expression of human cohesin in yeast could provide a research platform to investigate cohesin biology.

Human cohesin formed complexes and loaded onto chromatin resulting in dominant-negative effects in yeast. Human SMC1A and SMC3 subunits act like a dominant-negative subunit, disrupting the normal functions of yeast cohesin. Surprisingly, the effect was independent of ATP catabolism. Mutated human cohesin behaved similarly to wildtype human cohesin. By dissecting the various combinations of human and yeast cohesin subunits, it was apparent that while all human cohesin subunits could interact with yeast cohesin subunits, the dominant-negative phenotype was dependent on chromatin loading. For example, when expressed in the absence of human SMC proteins, the non-SMC subunits RAD21 and STAG2 co-immunoprecipitated with yeast Smc1 but did not associate with chromatin or cause a dominant-negative effect. The behaviour of human cohesin as a dominant-negative agent that was synthetic lethal in cells with mild cohesin defects raises the interesting possibility that cohesin could be targeted to elicit a dominant-negative effect that selectively kills cancer calls. Such a dominant synthetic lethal effect may be effective in targeting cancer cells that carry somatic mutations in a cohesin component or other cancer driver genes.

## MATERIALS AND METHODS

### Expression vectors

Human cohesin genes expressed individually were obtained as Gateway-compatible entry clones from the hORFeome V8.1 collection (Yang et al., 2011) and shuttled into yeast destination vectors pAG416GPD-ccdB+6Stop (*URA3*, CEN, constitutive GPD promoter, 6-amino-acid C-terminal extension) or pAG425GAL-hORF+6Stop (*LEU2*, 2µ, inducible GAL promoter, 6-amino-acid C-terminal extension) (Alberti et al., 2007; Kachroo et al., 2015) using LR Clonase II (Invitrogen) to generate expression clones. The neochromosomes yCL3A, hC, hCL, hCL2A, hCL6A, hL, h2A, hS/S and h3P (see Figure 1B) were synthesized by Neochromosome Inc. Neochromosome derivatives of hC with different deletions were constructed by CRISPR-mediated deletions using donor DNA generated from annealed oligos and sgRNAs targeted to the corresponding human ORFs. Neochromosome yCL3A-*smc1*Δ*smc3*Δ was constructed by CRISPR-mediated deletion of *SMC1* and *SMC3* using donor DNA generated from annealed oligos and sgRNAs targeted to linker sequences on the vector. GAL-inducible *URA3*-marked expression vectors of yeast cohesin genes were obtained from the FLEX collection (Hu et al., 2007). Yeast *SMC1* from the FLEX collection (*URA3*) was used to create a FLAG-tagged *SMC1* construct by inserting 3xFLAG at residue R373 in the coiled-coil domain via restriction cloning (donor tagged constructed gifted by the Vincent Guacci and Douglas Koshland) (Camdere et al. 2015). A Rad52-YFP fusion protein (C-terminally tagged) was expressed from the plasmid pRS413-RAD52-YFP (*HIS3*) from a RAD52 endogenous promoter (construct gifted by Rodney Rothstein) (Alvaro et al., 2006). For plasmid shuffle strains, yeast genes were amplified from genomic DNA and cloned to pRS416 (*URA3*), under the control of their endogenous promoter (pRS416-*RAD52*, pRS416-*CTF8*, pRS416-*RAD61*).

### Introducing mutations to neochromosomes

Mutational alterations to the neochromosomes were performed using CRISPR-Cas9 as a marker-less, precise editing system; described in this protocol (Quick and Easy CRISPR Engineering in Saccharomyces Cerevisiae · Benchling, by William Shaw). The sgRNA guides were designed using Benchling “Design and Analyze Guides”. The sgRNA were assembled into the sgRNA expression vector pWS082 using golden gate assembly. Donor DNA was constructed by annealing two complimentary oligos that contained both the intended missense mutation and a silent mutation to disrupt the PAM site. CRISPR components were transformed into yeast using a standard lithium acetate transformation protocol which included 100ng of linearized Cas9 vector, 200ng of digested sgRNA vector and ∼2μg annealed donor DNA. The transformations were plated on SC-Ura-Leu media to select for both the assembled sgRNA-Cas9 vector and the neochromosome. To confirm that the missense mutations were incorporated, transformants were streaked on SC-Leu media to allow loss of the Cas9-sgRNA vector but maintain the *LEU2*-marked neochromosome. To confirm correct editing, the neochromosomes were extracted from the yeast cells and sanger sequenced.

### Yeast strains

Expression vectors containing human or yeast cohesin genes and vector controls pRS415 (*LEU2*), pRS416 (*URA3*) or pRS413 (*HIS3*) (Sikorski and Hieter, 1989) were transformed into indicated yeast strains and maintained on selective media. Yeast strains included wildtype *MAT*α BY4742 (Brachmann et al., 1998), conditional temperature-sensitive mutants of cohesin genes (Li et al., 2011), *rad61*Δ, *ctf8*Δ, and *rad52*Δ which were generated by sporulating heterozygous deletion strains (Giaever et al., 2002) and yeast cohesin complex deletion strains (ΔC, ΔCL, ΔCL2A and ΔCL3A). To create the multi-subunit yeast cohesin deletion strains (4, 6, 8, and 9 gene deletions), donor DNA generated by PCR was co-transformed into a yCL3A-containing BY4742 strain along with linear fragments encoding Cas9 and sgRNAs targeted to the unique endogenous terminators of the yeast genes (except for *SCC4*, which targeted the yeast ORF). Gene deletions were completed sequentially starting with the 4 core cohesin genes (ΔC to ΔCL3A) and confirmed by PCR. A range of Cas9-sgRNA expression vectors with differing selection markers were utilized for the iterative gene deletions such that once a transformant was confirmed for a single deletion, a new round of transformation was started without the need to rid the cells of the previous CRISPR/Cas9 machinery. Donor DNA was obtained from PCR using yeast genomic DNA as template and primers were designed such that one primer contained (5’ to 3’) ∼60bp homology to the region upstream of the yeast ORF and ∼25bp homology to the endogenous terminator of the yeast gene, while the other primer was designed to be homologous to the endogenous terminator in a region downstream of the corresponding primer pair.

### Plasmid shuffle assay to assess complementation and synthetic lethal interactions

To assess complementation of cohesin complex deletion mutants, yeast strains were grown to saturation in SC−Leu−His at 30°C to allow loss of *URA3*-marked yCL3A. Approximately 10^7^ cells were then plated on SC−Leu−His and SC−Leu−His+5-FOA (0.1%) to select for Ura- segregants via the plasmid shuffle strategy (Boeke et al., 1987) and incubated at 30°C. Plates containing 5-FOA were incubated for a minimum of 10 days and any viable colonies were streaked on new 5-FOA plates before isolation of vector DNA. To confirm presence/absence of *URA3*-marked yCL3A, two sites corresponding to the *URA3* gene and a yeast cohesin gene on the vector were chosen as template for PCR. In all cases tested, 5-FOA resistant colonies were confirmed to contain yCL3A and viability of the cohesin deletion mutants was attributed to the presence of the yeast cohesin genes on yCL3A. Presumably, the 5-FOAr (Ura-) phenotype was due to mutations in the *URA3* gene present on yCL3A. The plasmid shuffle method was also used to test for dominant synthetic lethality caused by human proteins expressed from a human neochromosome in strains carrying genomic deletions of *rad52Δ, ctf8Δ* and *rad61Δ;* in these cases the strains carried *URA3*-marked plasmids carrying the respective *RAD52*, C*TF8* or *RAD61* genes to allow uncovering of the corresponding genomic deletion in 5-FOA media.

### Growth assays

Yeast serial spot dilutions were performed as a semi-quantitative growth assay. Strains were grown overnight in selective media then 5μl were spotted in 10x serial dilutions. Plates were grown 2-5 days at 25°C, 30°C or 37°C depending on the experiment. For liquid growth assays, cultures were grown to mid-log phase then diluted to OD_600_=0.1 in 200µl media +/- 0.0075% MMS. OD_600_ readings were measured every 30 minutes over a period of 24h in a TECAN M200 plate reader and plates were shaken for 10 minutes before each reading. Strains were tested in 3 replicates per plate per condition and area under the curve (AUC) was calculated for each replicate. Strain fitness was defined as the AUC of each yeast strain relative to the AUC of the control strain (BY4742 + vector controls) grown on the same plate in the same media condition. For induction in galactose media, yeast strains were grown to mid-log phase in both dextrose or galactose media before diluting to OD_600_=0.1 in the same media +/-0.0075% MMS. All liquid growth assays were done at 30°C.

### Rad52-YFP foci to quantify homologous recombination intermediates

Rad52-YFP foci quantification was used to evaluate levels of homologous recombination in different conditions. Wildtype strains containing the hC or hCL neochromosomes were transformed with the pRS413-*RAD52*-YFP plasmid and incubated at 25°C. Transformants were grown in liquid double selection media (SC−Leu−His) overnight. Cells were diluted to an OD_600_ of 0.3 and grown for 4 hours to logarithmic growth phase. After seeding onto a Concanavalin A treated slide, YFP foci were imaged using a Zeiss confocal fluorescence microscope. Images were taken with 6 Z-stacks and maximum intensity projection images were generated. At least 100 cells per image were counted per condition and compared to the number of Rad52-YFP foci percentage per condition.

### Chromatin fractionation to assess human cohesin subunits interacting with yeast chromatin

Chromatin fractionation was performed using a sucrose gradient method. Strains were grown overnight in selective media then diluted to OD_600_ 0.2 in a 50mL liquid culture and grown until OD_600_ ∼1. Cells were harvested by centrifugation and washed once with dH_2_O and once with SB Buffer (1M Sorbitol, 20mM Tris Cl pH 7.4). Cell pellets were stored at -80°C until use. To fractionate the cells, the pellet was thawed on ice and subsequently washed with 1mL 1x PBS (+10mM β-Me, added fresh), followed by a wash with 1mL SB buffer. Cells were resuspended in 1mL SB and transferred to a 2mL tube. To digest the cell wall, a final concentration of 1.25mg/mL Zymolase-20T (from *A. Luteus*)(Seikagaku BioBusiness Cat. No. 120491) was added and cells were rotated at room temperature. Digestion was monitored by OD_600_ and continued until >80% of cell walls were digested, approximately 1 hour. Spheroplasted cells were then centrifuged at 2500RPM for 5 minutes and gently washed with 1mL SB. Spheroplasts were then resuspended in 500μL EBX buffer (20mM Tris Cl pH 7.4, 100mM NaCl, 0.25% Triton X-100) with protease inhibitors (1mM PMSF, 1μM Leupeptin, 1μM Pepstatin A, 2μg/mL Aprotinin), 15mM β-Me and 0.5% Triton X-100, added fresh, then set on ice to lyse the outer membranes for 10 minutes with gentle inversions every 2 minutes. In a new 2mL tube, 1mL of NIB buffer (20mM Tris Cl pH 7.4, 100mM NaCl, 1.2M Sucrose) with protease inhibitors (1mM PMSF, 1μM Leupeptin, 1μM Pepstatin A, 2μg/mL Aprotinin) and 15mM β-Me, added fresh, was added to the tube and the lysates were gently layered over top. The samples were then centrifuged at 4°C, 14000 RPM for 15 minutes to separate the nuclear pellet from the cytoplasmic proteins. 100μl was then pipetted from the top layer and saved as the cytoplasmic fraction. The remaining supernatant was removed and the nuclei (pellet) were resuspended in 500ml EBX buffer with protease inhibitors, 15mM β-Me and 1% Triton X-100. This mixture was incubated on ice for 10 minutes and inverted every 2 minutes to allow for nuclear lysis. The samples were then centrifuged for 10 minutes at 4°C, 14000 RPM. The resulting pellet (chromatin) was washed once with 500μl EBX and centrifuged again for 10 minutes at 4°C, 14000 RPM. The chromatin pellet was resuspended in 100μl Tris HCL pH 8.0 and saved as the chromatin fraction. When applied to Western Blotting, SDS buffer (62.5mM Tris HCL pH 7.5, 8% SDS, 10% Glycerol, 0.005% Bromophenol blue + 355mM β-Me added fresh) was added, samples were boiled for 5 minutes and centrifuged at 9000 RPM for 2 minutes. 10μl of cytoplasmic fraction and 2μl of chromatin fraction were run on the gel, to account for differences in protein density.

### Co-immunoprecipitation between yeast and human cohesin subunits

Co-IP was performed using Invitrogen Protein A Dynabeads (Invitrogen, Cat No. 10001D) and performed as described by the manufacturer’s instructions. To extract proteins for Co-IP, yeast strains were grown overnight in selective media. Saturated cultures were diluted to OD_600_ ∼0.2 in a 50mL liquid culture and grown until OD_600_ ∼1, at 30°C. Cells were harvested by centrifugation, washed with dH_2_O and pellets were stored at –80°C until use. To perform the IP, cell pellets were thawed on ice and resuspended in 0.5mL Lysis buffer (50mM NaCl, 1mM EDTA, 50mM Hepes pH 7.4) with 0.1% Tween and freshly-added protease inhibitors (1mM PMSF, 1μM Leupeptin, 1μM Pepstatin A, 2μg/mL Aprotinin), and incubated for 10 minutes on ice to allow for lysis of the cells. About 100μl of 0.5mm diameter glass beads (BioSpec Products, Cat No. 11079105) were added to the lysates. Samples were vortexed for 45 seconds and then placed on ice for 5 minutes. This was repeated 3 times and samples were then centrifuged at 10000 RPM for 10 minutes at 4°C. Concurrently, 30μl of Dynabeads Protein A were prepared per sample, as described by the manufacturer (conjugated with antibody at 10μg/30μl beads). Following centrifugation, 50μl of the supernatant was saved as the pre-IP (inputs), mixed with equal parts 2x Laemmli Sample Buffer (4% SDS, 20% Glycerol, 120mM Tris-HCl pH 6.8, bromophenol blue + 10% β-Me (added fresh)) and frozen at -80°C for storage. The remaining supernatant (∼450μl) was transferred onto the pre-conjugated Dynabeads. Samples were rotated at room temperature for 60 minutes. The beads were then washed 3x with 500μl Lysis Buffer (+PIs, +Tween) by vortexing for 10 seconds each time. The samples were then eluted with 50μl Thorner’s Buffer (5% SDS, 3.8M Urea, 40mM Tris pH 6.8, 0.1mM EDTA) + 1% β-Me (added fresh) by incubating at 80°C for 5 minutes. When visualizing by Western Blot, 5μl of both the inputs and IPs were loaded onto the gel.

### Western Blotting

Western Blotting was used to visualize proteins in the chromatin fractionation and Co-IPs. 10% or 12% polyacrylamide gels were made for Western blotting (BioRad Mini-PROTEAN Handcast system, Cat. No. 1658001) and proteins were transferred to pre-activated PVDF membranes (Millipore, IPVH00010). Membranes were imaged using a BioRad ChemiDoc transilluminator. The following antibodies were used in this study SMC1A (1:1000, Abcam ab9262), SMC3 (1:1000, Abcam ab9263), RAD21 (1:1000, Millipore 05-908), STAG2 (1:1000, Bethyl A302-581), FLAG M2 (1:3000, Sigma F3165), Pgk1 (1:1000, Santa Cruz sc-130335), Cpy (1:5000, Abcam ab113685) and Acetyl H4 (1:10000, Sigma 06-866).

### Single-molecule sequencing of co-immunoprecipitated proteins on a Quantum SI Platinum machine

After co-immunoprecipitation in 50 μL of buffer, the sample was split with 15 μL run on a Western blot and 30 μL used to make a peptide library for sequencing as described in the Platinum® Library Preparation Kit V2 Protocol. Briefly, the magnetic beads were pelleted and the co-immunoprecipitation buffer was replaced with 100 μL of QSI Sample Buffer. Reduction, alkylation, and Lys-C digestion steps were performed with no centrifugation. Peptides were activated and 2 μL CTAB, 2 μL EDTA and 1 μL of K-linker from the library preparation kit were added. SMPS was performed using the V4 Platinum Sequencing Kit following the recommended protocol. Data were analyzed with the Primary Analysis V2.16.0 algorithm, and peptides were aligned to reference libraries containing yeast and human cohesin protein sequences using the Peptide Alignment V18.2 algorithm. Sequenced peptides that aligned to STAG2 (amino acids 785-854) were excluded from analysis as it shared a recognizer motif with the repeat segment of Protein A, which was present on the magnetic beads used to pulldown the cohesin complexes. As a result this peptide was highly abundant in the sequenced sample.

### FACS in yeast to analyze cell cycle progressions

Cell cycle analysis was performed with logarithmically growing yeast cells to assess cell cycle progression (as described in Oppedisano et al. 2025). 1mL of logarithmic phase cells were then collected for analysis. Thirty thousand cells were counted per condition using a CytoFlex machine.

### Modelling 3D structures of cohesin

All SMC structures were generated using Alphafold3 and were visualized with ChimeraX10 software (Abramson et al., 2024; Meng et al., 2023). Yeast-human hybrid cohesin models were generated by inputting different combinations of yeast proteins (Smc1, Smc3, Irr1, Mcd1) using sequences from Saccharomyces Genome Database (SGD) (YFL008W, YJL074C, YIL026C, YDL003W, respectively) and human proteins (SMC1A, SMC3, STAG2 and RAD21) using sequences from UniProt (Q14683, Q9UQE7, Q8N3U4, Q3SWX9 respectively).

## FUNDING

This work was funded by a grant from CIHR to PCS (project 486753).

## DATA AVILABILITY

All data necessary to reach our conclusions are included within the manuscript and Supplemental Figures. All strains, plasmids, neochromosomes and other resources are available through contacting the corresponding authors.

## Supplemental

**SUPPLEMENTAL FIGURE 1:**
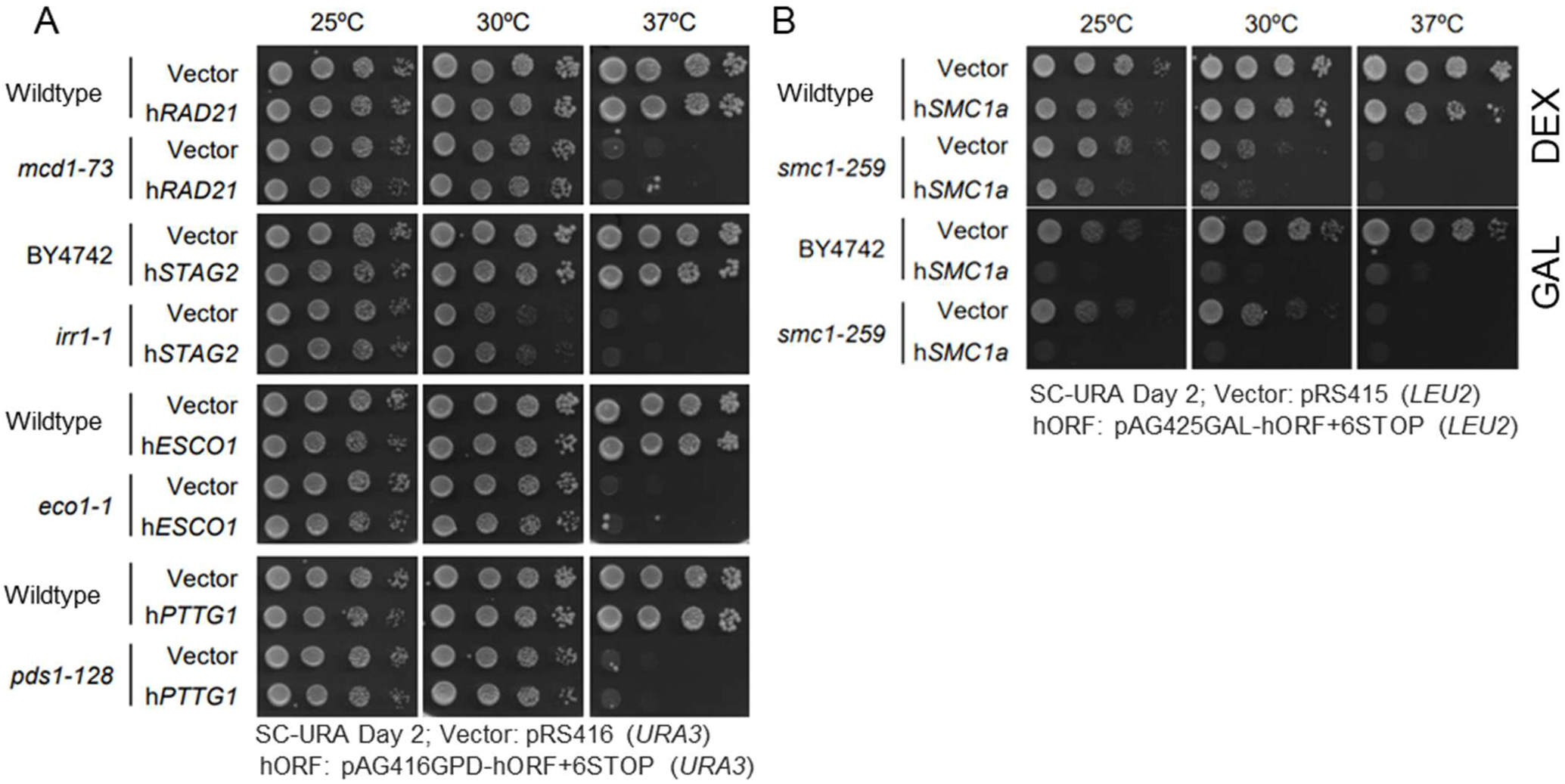
Human cohesin subunits do not complement yeast cohesin subunits when expressed individually. (**A**) Human cohesin subunits, being expressed from a GPD promoter, in wildtype and corresponding temperature sensitive strains (*RAD21* + *mcd1-73*, *STAG2* + *irr1-1*, *ESCO1* + *eco1-1*, *PTTG1* + *pds1-128*). Serial spot dilution plates were incubated at restrictive temperatures to test complementation. (**B**) Galactose-inducible expression of *SMC1A* in wildtype yeast causes a dominant-negative phenotype at permissive temperatures and does not complement the *smc1-259* mutation at the restrictive temperature.

**SUPPLEMENTAL FIGURE 2:**
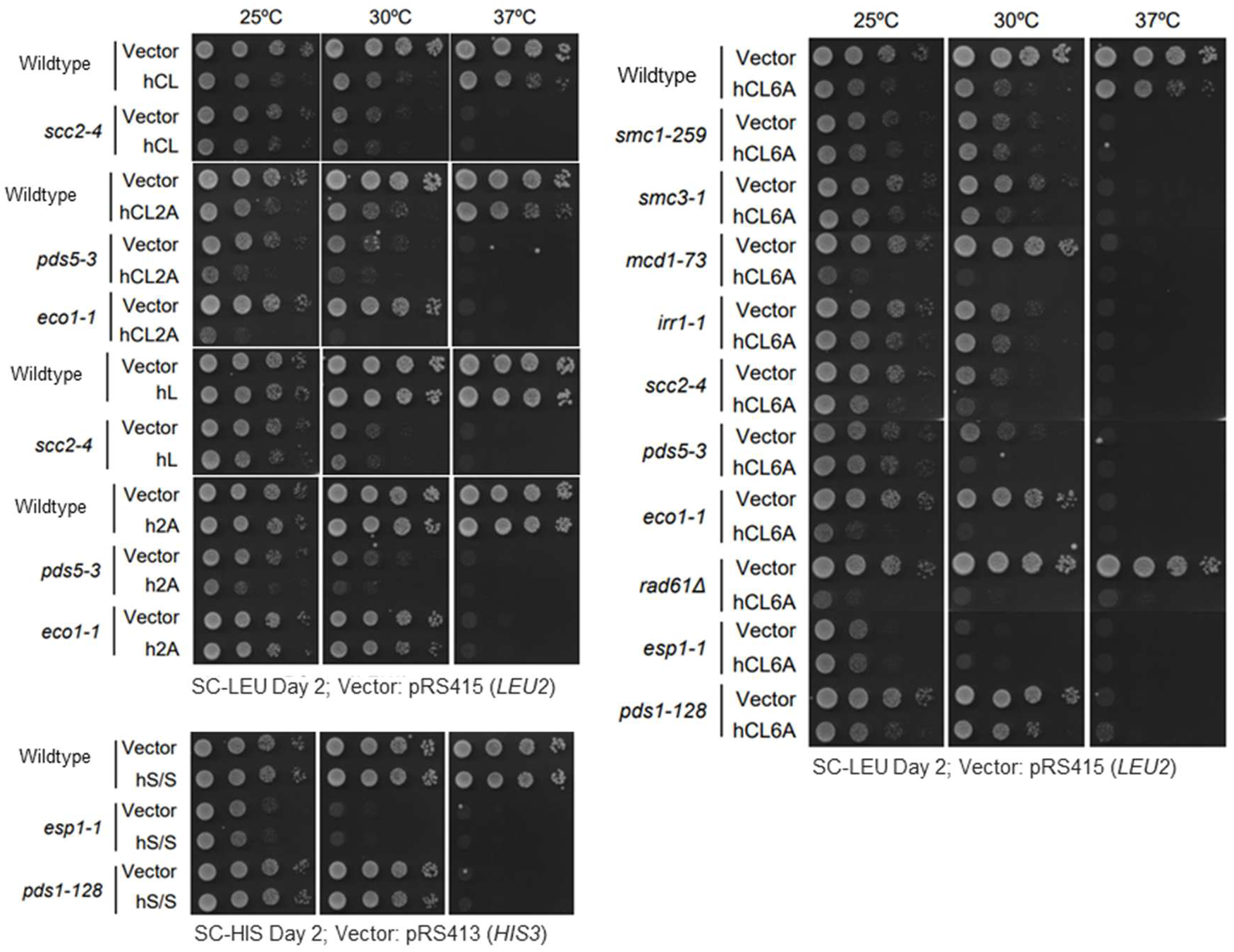
Different variations of human cohesin neochromosomes fail to rescue temperature-sensitivity of single yeast cohesin temperature-sensitive mutants. This includes hCL (*scc2-4*), hCL2A (*pds5-3, eco1-1*), hL (*scc2-4*), h2A (*pds5-3, eco1-1)* and hS/S (*esp1-1, pds1-128*). Further, hCL6A was tested for complementation for all corresponding yeast genes (*smc1-259, smc3-1, mcd1-73, irr1-1, scc2-4, pds5-3, eco1-1, eps1-1* and *pds1-128*). Notably, human cohesin neochromosomes that contain human core proteins caused dominant negative fitness defects in wildtype yeast, and in some cohesin mutant backgrounds, this effect was amplified.

**SUPPLEMENTAL FIGURE 3:**
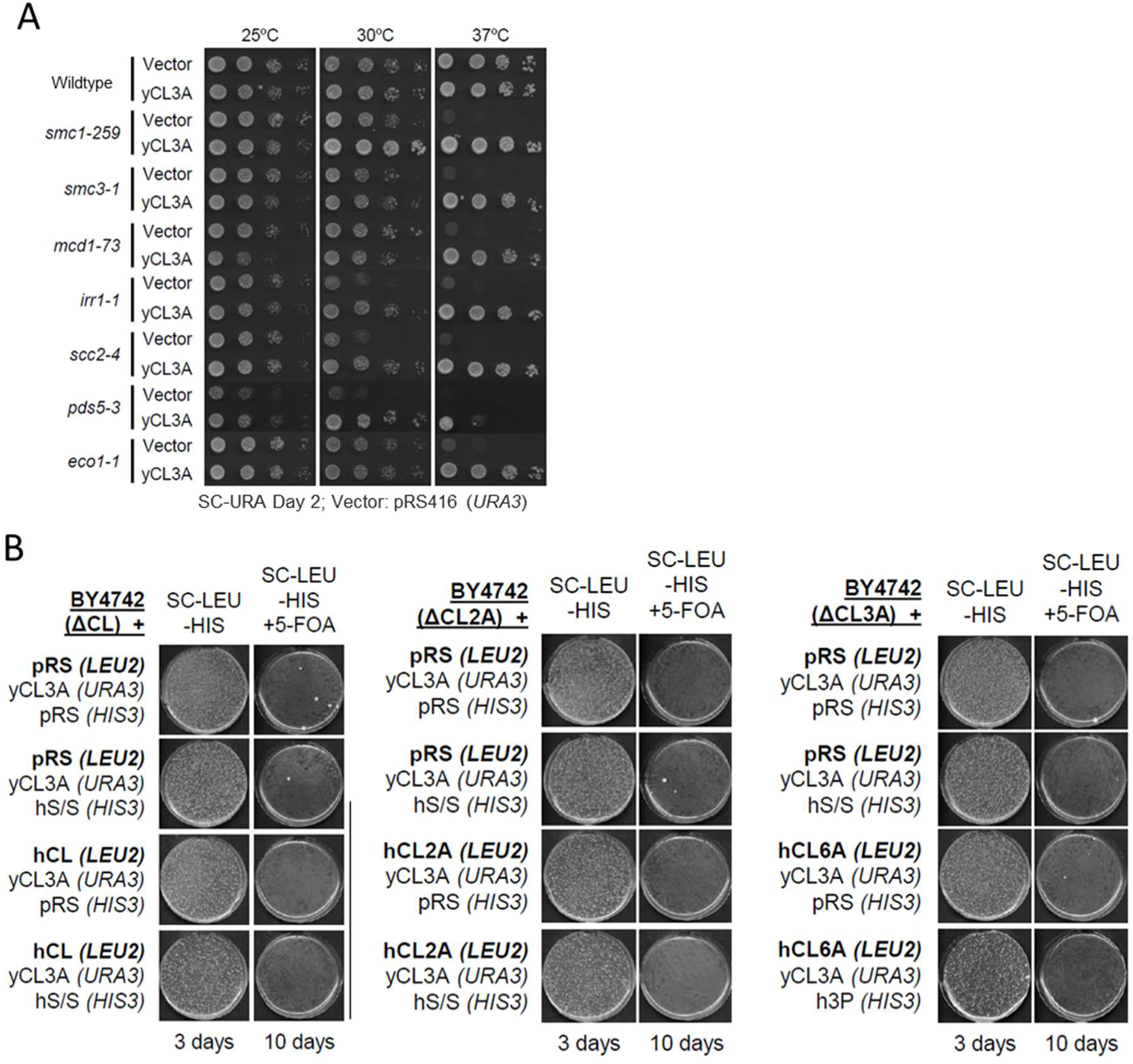
Simultaneously replacing up to nine cohesin yeast genes with the human orthologs does not support viability. (**A**) Yeast cohesin genes expressed from yCL3A (*SMC1*, *SMC3*, *MCD1*, *IRR1*, *SCC2*, *SCC4*, *PDS5*, *RAD61*, *ECO1*) functionally complement yeast loss of function mutations at the restrictive temperature. (**B**) Strains with yeast cohesin genes genomically deleted (ΔCL: *smc1Δ, smc3Δ, mcd1Δ, irr1Δ scc2Δ scc4Δ*; ΔCL2A: *smc1Δ, smc3Δ, mcd1Δ, irr1Δ scc2Δ scc4Δ pds5Δ eco1Δ*; ΔCL3A: *smc1Δ, smc3Δ, mcd1Δ, irr1Δ scc2Δ scc4Δ pds5Δ eco1Δ rad61Δ*) expressing yCL3A to complement genetic knockouts, and transformed with equivalent human neochromosomes (hCL, hCL2A, or hCL6A). When plated of 5-FOA, viability was not supported by human cohesin genes (+/- hS/S or h3P).

**SUPPLEMENTAL FIGURE 4:**
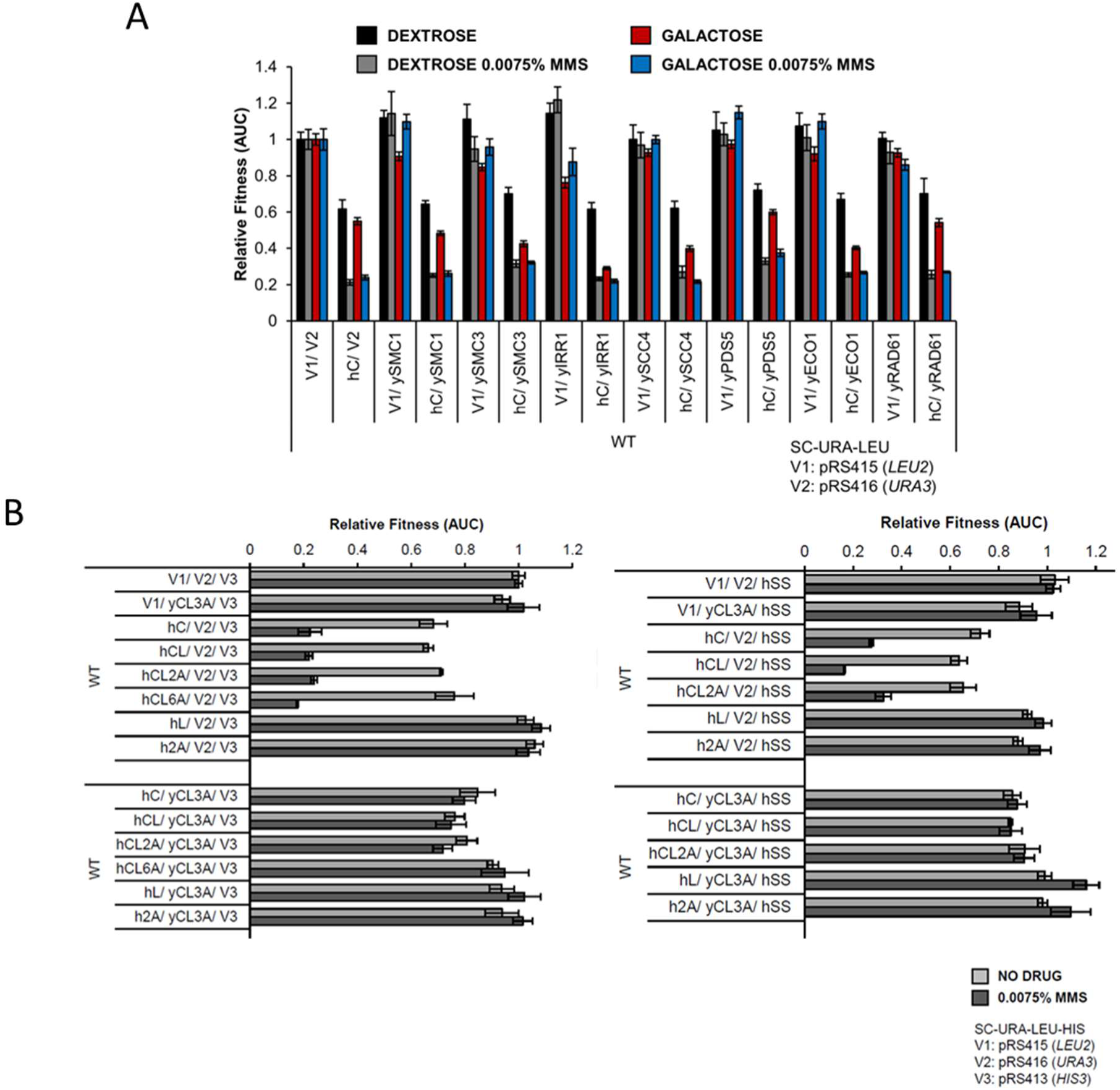
Dominant negative phenotype of hC can be suppressed with yCL3A, but not individual yeast subunits. (A) Inducible overexpression of individual yeast cohesin subunits does not rescue the dominant negative growth phenotype caused by expression of hC, in the presence or absence of MMS. (B) Co-expression of yCL3A with either hC, hCL, hCL2A, hCL6A, hL or h2A (without hS/S (left panel) and with hS/S (right panel) in wildtype cells results in a rescue of the growth phenotypes caused by the presence of the human core subunits.

**SUPPLEMENTAL FIGURE 5:**
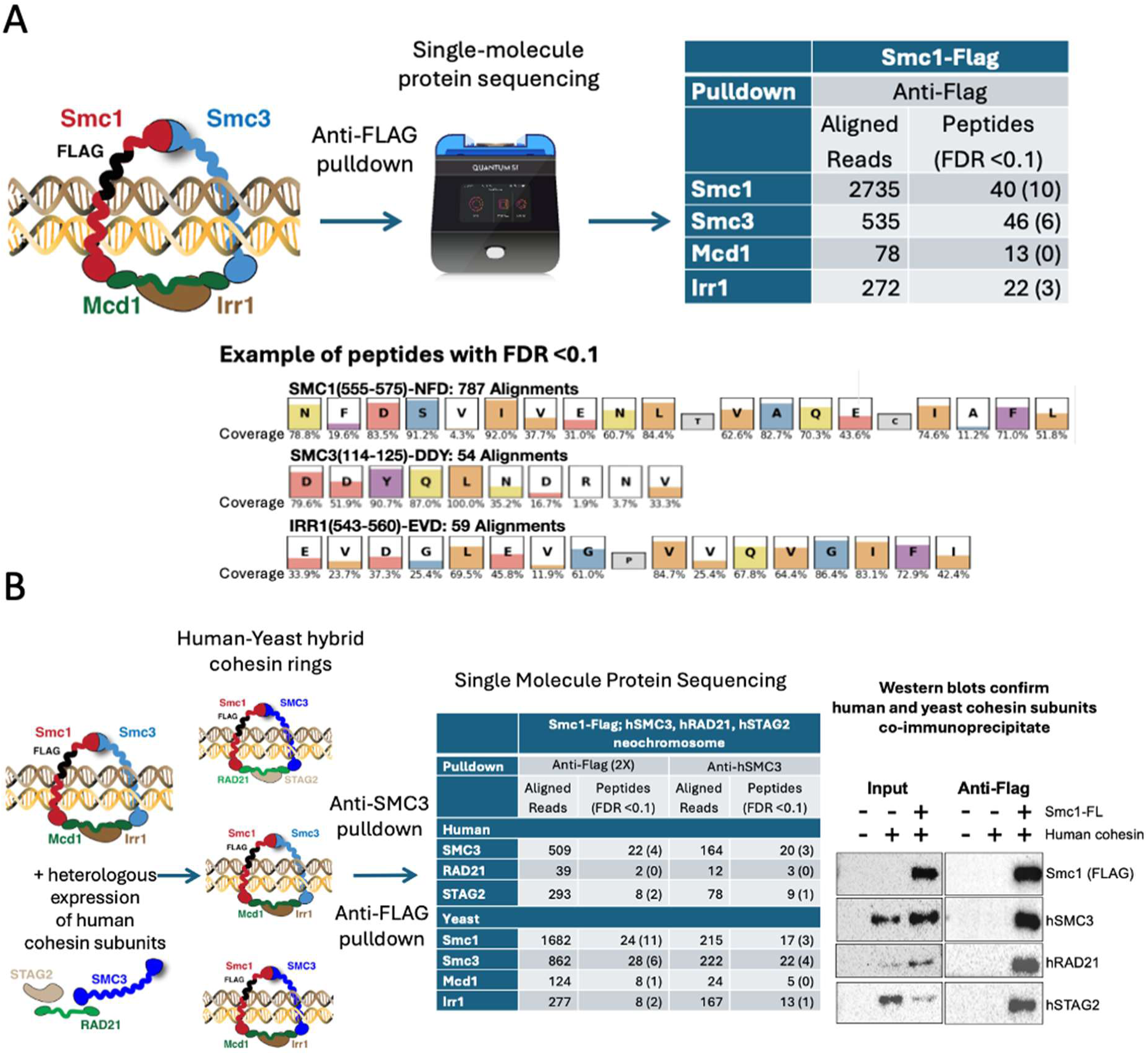
Single-molecule protein Sequencing of cohesin complexes immunoprecipitated from yeast cells. (A) SMPS identifies yeast cohesin subunit peptides from samples co-immunoprecipitated with an anti-FLAG antibody from yeast cells expressing FLAG-Smc1. Representative data of Smc1, Smc3, and Irr1 peptides identified with a false discovery rate (FDR) <0.1. (B) SMPS identifies both yeast and human cohesin subunit peptides from samples co-immunoprecipitated with an anti-Flag antibody from yeast cells expressing FLAG-Smc1 and human SMC3, STAG2 and RAD21. Western blotting confirms the presence of yeast FLAG-Smc1, human SMC3, human RAD21 and human STAG2 in the immunoprecipitated pellet.

**SUPPLEMENTAL TABLE 1:** Summary of complementation efforts with neochromosomes, in Ts strains and as plasmid shuffle swaps.

|  | Yeast strain | hC | hCL | hCL2A | hCL6A | hL | h2A | hS/S |
| --- | --- | --- | --- | --- | --- | --- | --- | --- |
| **Core complex** | *smc1-259* | NO |  |  | NO |  |  |  |
|  | *smc3-42* | NO |  |  | NO |  |  |  |
|  | *mcd1-73* | NO |  |  | NO |  |  |  |
|  | *irr1-1* | NO |  |  | NO |  |  |  |
| **Loader Complex** | *scc2-4* |  | NO |  | NO | NO |  |  |
| **Accessories** | *pds5-3* |  |  | NO | NO |  | NO |  |
|  | *eco1-1* |  |  | NO | NO |  | NO |  |
|  | *rad61Δ* |  |  |  | NO |  |  |  |
|  | *esp1-1* |  |  |  | NO |  |  | NO |
|  | *pds1-128* |  |  |  | NO |  |  | NO |

|  | hC | hCL | hCL2A | hCL6A | hC hS/S | hCL hS/S | hCL2A hS/S | hCL6A h3P |
| --- | --- | --- | --- | --- | --- | --- | --- | --- |
| **ΔC:** *smc1Δ smc3Δ mcd1Δ irr1Δ* | NO |  |  |  | NO |  |  |  |
| **ΔCL:** *smc1Δ smc3Δ mcd1Δ irr1Δ scc2Δ scc4Δ* |  | NO |  |  |  | NO |  |  |
| **ΔCL2A:** *smc1Δ smc3Δ mcd1Δ irr1Δ scc2Δ scc4Δ pds5Δ eco1Δ* |  |  | NO |  |  |  | NO |  |
| **ΔCL3A:** *smc1Δ smc3Δ mcd1Δ irr1Δ scc2Δ scc4Δ pds5Δ eco1Δ rad61Δ* |  |  |  | NO |  |  |  | NO |

